# Abundant microorganisms in soil are mainly controlled by stochastic processes, while bacteria and fungi differ in rare taxa

**DOI:** 10.1101/2023.12.14.571670

**Authors:** Guo Qian, Lu Gong

## Abstract

The assembly mode of microbial communities helps to explain the ecological processes of soil subsurface groups, and abundant and rare microorganisms have their own unique assembly patterns. In this paper, the effect of changing vegetation types on the assembly of bacteria and fungi with different abundances in soil was investigated by iCAMP in the Tianshan Mountains. The results showed that: (i) there were differences in the distribution of rare and abundant microbial communities among different vegetation types: the main trends were forests>shrubs > grasslands, and the α diversity and variability of rare microorganisms were greater than those of abundant taxa; (ii) abundant taxa were mainly affected by stochastic processes (mainly diffusion limitation), whereas among rare microbes, the main controlling process for bacteria was heterogeneous selection and for fungi was diffusion limitation; (ⅲ)soil carbon, nitrogen and temperature were important determinants driving bacterial community structure. Our results deepen the understanding of the various ecological processes involved in microbial community assembly, reveal the effects of environmental factors on abundant and rare microorganisms, and provide evidence for understanding the mechanisms of soil microbial community construction among different vegetation covers in arid zones.

**Importance:** The study compares the response of different vegetation types to soil microbial community processes in the arid zone and deepens the understanding of the mechanisms of soil microbial community composition at different abundances.

## 1. Introduction

The distribution of soil microbial community multiplicity tends to be uneven, dominated by a few abundant species, while a large number of other species are in low abundance [1]. Rarity and abundance of microbial richness exhibit quite different biogeographic patterns due to differences in competition, dispersal and resilience, among others. Enriched microorganisms effectively utilise a wider range of resources than rare taxa[2], have wider ecological niche thresholds, and are more tolerant and resilient to harsh environments[3]. The relative abundance of microbial taxa may play an important role in regulating the assembly of different communities, e.g., slight abundance changes due to ecological drift can significantly affect low-abundance microbial populations [4]. Recent studies have begun to explore the relationship between community assembly processes and the relative abundance of microbial taxa by dividing microbial communities into two sub-communities based on relative abundance thresholds [5, 6], but studies in dryland forests remain rare.

Previous studies have generally divided the biome building process into two categories: deterministic and stochastic processes, which stem from two different theories. The neutral theory holds that all species have equal ecological functions and that species dynamics are controlled by stochastic processes[7]. Niche-based theories suggest that non-random and niche-based ecological processes control diversity patterns, and these are known as deterministic processes[8]. In reality, these two processes often coexist, and the assembly process of microbial communities can be characterized by quantifying the degree of action of the two ecological processes. The previous researchers used correlation or heatmap method to present the community construction, but due to the complexity of microbial species and branches, the correlation analysis alone would discard the information or fail to present the whole phylogeny; Ning, Yuan [9]proposed a framework to quantitatively infer the mechanism of community assembly based on phylogenetic zero model analysis (iCAMP), which divides the community aggregation process into five categories: homogeneous selection (HoS), heterogeneous selection (HeS), homogeneous diffusion (HD), diffusion restriction (DL), and drift (DR). HeS and HoS are deterministic processes that emphasize the role of the environment and consider abiotic factors that prevent species from establishing or persisting in local communities as “environmental filters”[10]. Based on a null model and phylogenetic signals reflecting ecological niche differences within certain thresholds, the framework analyses the mechanisms of community construction at the level of the phylogenetic bin, quantifying the assembly process at the level of individual taxa/spectra rather than the entire community, thus enabling a mechanistic understanding of community diversity and succession that shows a high degree of accuracy.

The relative importance of deterministic and stochastic processes for the assembly of soil microbial communities varies according to environmental gradients (e.g. pH, nutrients, and climate) and the composition of different vegetation. In ecosystems with relatively homogeneous environments, stochasticity may be crucial [11], whereas in non-homogeneous environments with strong selectivity, deterministic processes tend to dominate, i.e., the “environmental filter” effect. Similarly, the presence of dominant species in different vegetation types generates “biological filters”, where the diversity of inter- and intraspecific interactions determines the assemblage of species in the local community and thus the ecological niche they achieve. Changes in the degree of dispersal of forest soil communities may be related to the diversity of species within habitats, where plant diversity promotes environmental homogeneity by reducing the complexity of plant-soil interactions and facilitating community convergence, thereby increasing stochasticity. At the same time, changes in the soil environment, including nutrient inputs and hydrothermal conditions, lead to changes in species rarity and ecological building processes.

Located in the hinterland of Central Asia, the TianShan Mountain of Xinjiang is rich in species resources, but quantitative studies on the ecological processes of microbial communities are still insufficient, especially in natural forest stands. In this paper, we used amplicon sequencing of 16S rRNA (indicative of bacterial communities) and ITS (indicative of fungal communities) to characterise microbial communities, and investigated the composition and construction mechanisms of microbial communities with different abundances of bacteria and fungi under forest soils in the Tian Shan. The aim of this study was to determine (1) the diversity patterns of microbial communities under different covers; (2) to quantify the relative importance of microbial assembly processes under different vegetation covers using the “iCAMP” framework, and the effects of community assembly processes on microbes of different abundances; (3) to analyse the extent to which soil factors influence different ecological processes, and explore the potential driving factors, in order to provide support for the study of forest microbial community assembly in arid zones.

## 2. Result

### 2.1 Taxonomic composition of abundant and rare fungi in different vegetation types

The Shannon index was used to characterise the α-diversity of microorganisms with different abundances in the soil, and there were significant differences in the diversity of bacterial and fungal communities among the five vegetation types (p < 0.001, Fig. 1), with both bacteria and fungi showing higher diversity in rare taxa. Specifically, abundant bacteria were found in mixed forests (4.80) > pure forests (coniferous forests: 4.78, broad-leaved forests: 4.73) > shrubs (4.70), and grasses (4.69), and rare bacteria were found in mixed forests (7.18), shrubs (7.13) > pure forests (coniferous forests: 7.06, broad-leaved forests: 7.03), and grasses (7.00), and the degree of variability was higher for rare taxa among the ecosystems. The degree of variation was greater for rare taxa in all cases. Fungi showed a trend similar to bacteria, but the variation was greater than that of bacteria and the regularity was not obvious, mainly showing that forests were greater than herbs. Among the different sampling sites, the highest α-diversity of abundant microorganisms was found in the ZTS, and the highest point of rare microorganisms was found in the XTS.

**Fig. 1.**
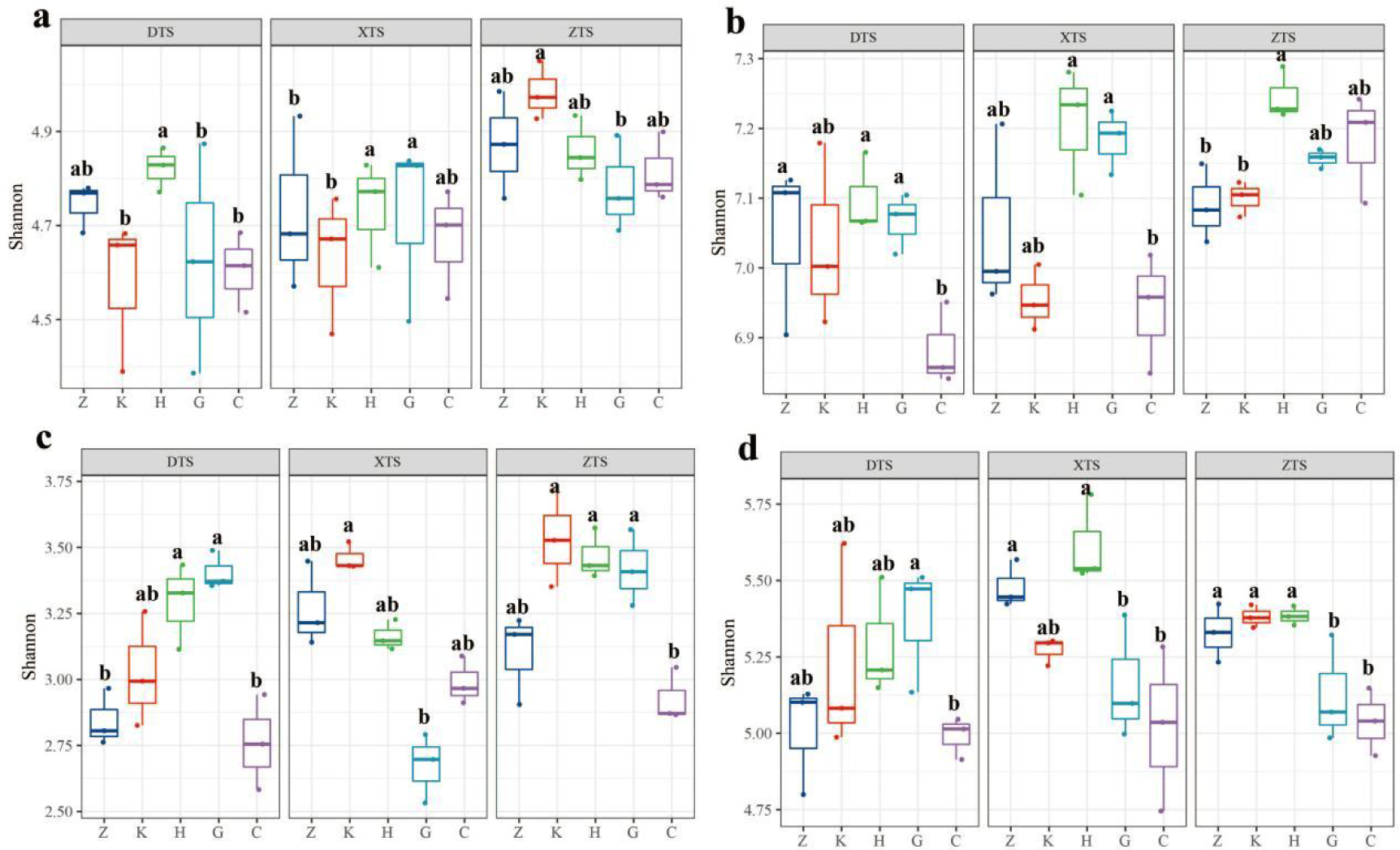
Alpha diversity of bacteria (a: abundant taxa, b: rare taxa)and fungi (c: abundant taxa, d: rare taxa)

Clustering and abundance analyses of the samples, the results of which are shown in Figure 2, indicate that the classification of fungi and bacteria with different abundances at the phylum level is virtually fixed, with differences in abundance. Of the total bacterial OTUs, 1.86% (239) were classified as abundant taxa and 98.14% (12,650) as rare taxa; of the total number of fungal OTUs, 4.88% (339) were abundant taxa and 95.12% (6,612) were rare taxa. The relative abundance results indicated that *Proteobacteria*, *Actinobacteria* and *Acidobacteriota* were the main phylum of bacteria, while the main body of the fungal community consisted of *Ascomycota*, *Basidiomycota*, *Chytridiomycota*, and *Mortierellomycota*. Among the abundant bacterial subcommunities, *Proteobacteria* (22.85%) were the most abundant and significantly higher in herbaceous types than in other types, and among rare bacteria, the distribution was more even among the types. Among the fungal communities, *Ascomycota* was overwhelmingly dominant and most abundant in the shrub types, and the taxonomic diversity of the abundant taxa was significantly lower than that of the rare taxa.

**Fig. 2.**
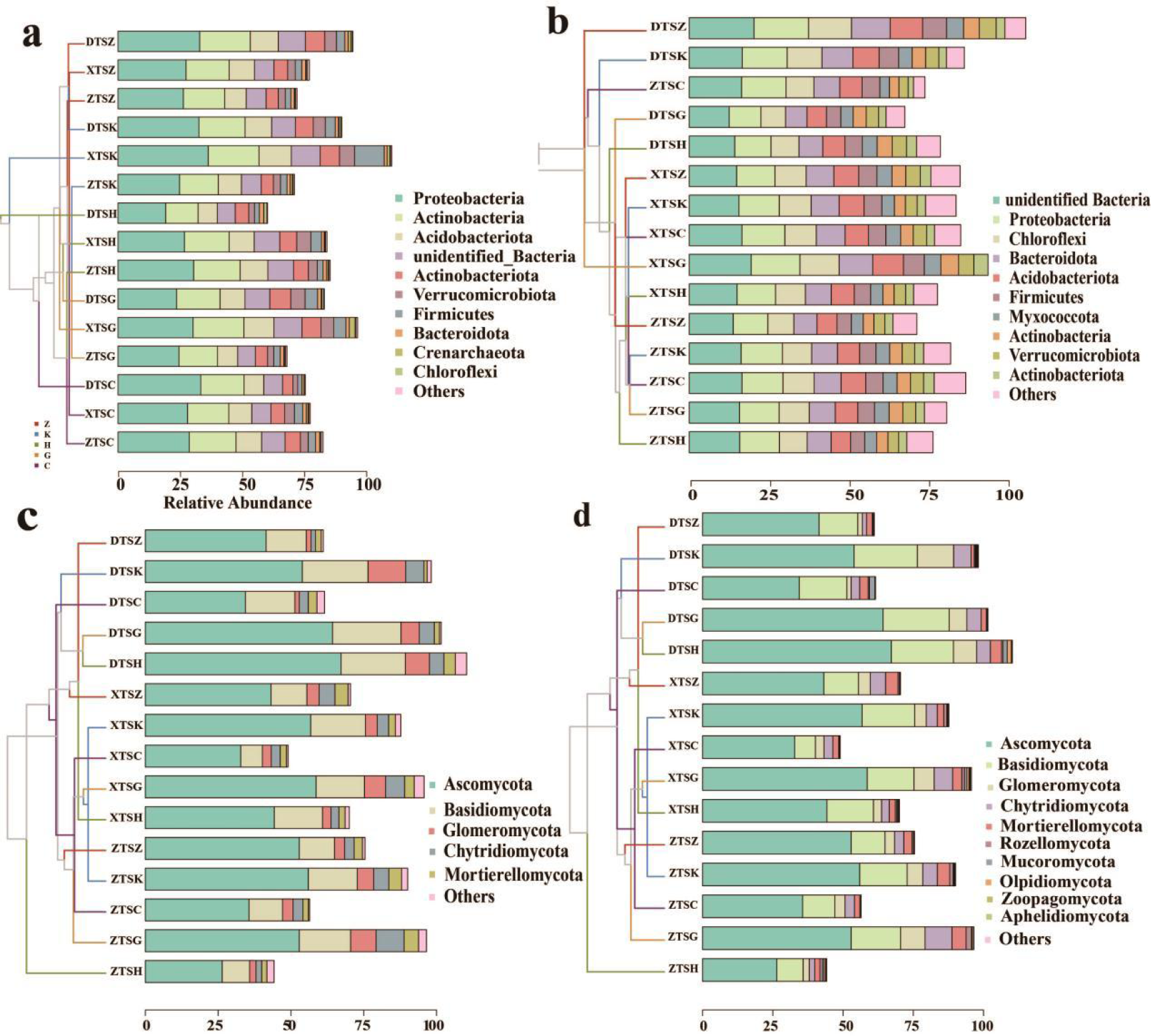
Distribution of microorganisms and clustering of samples (a: abundant bacteria, b: rare bacteria, c: abundant fungi, d: rare fungi)

**Figure 3.**
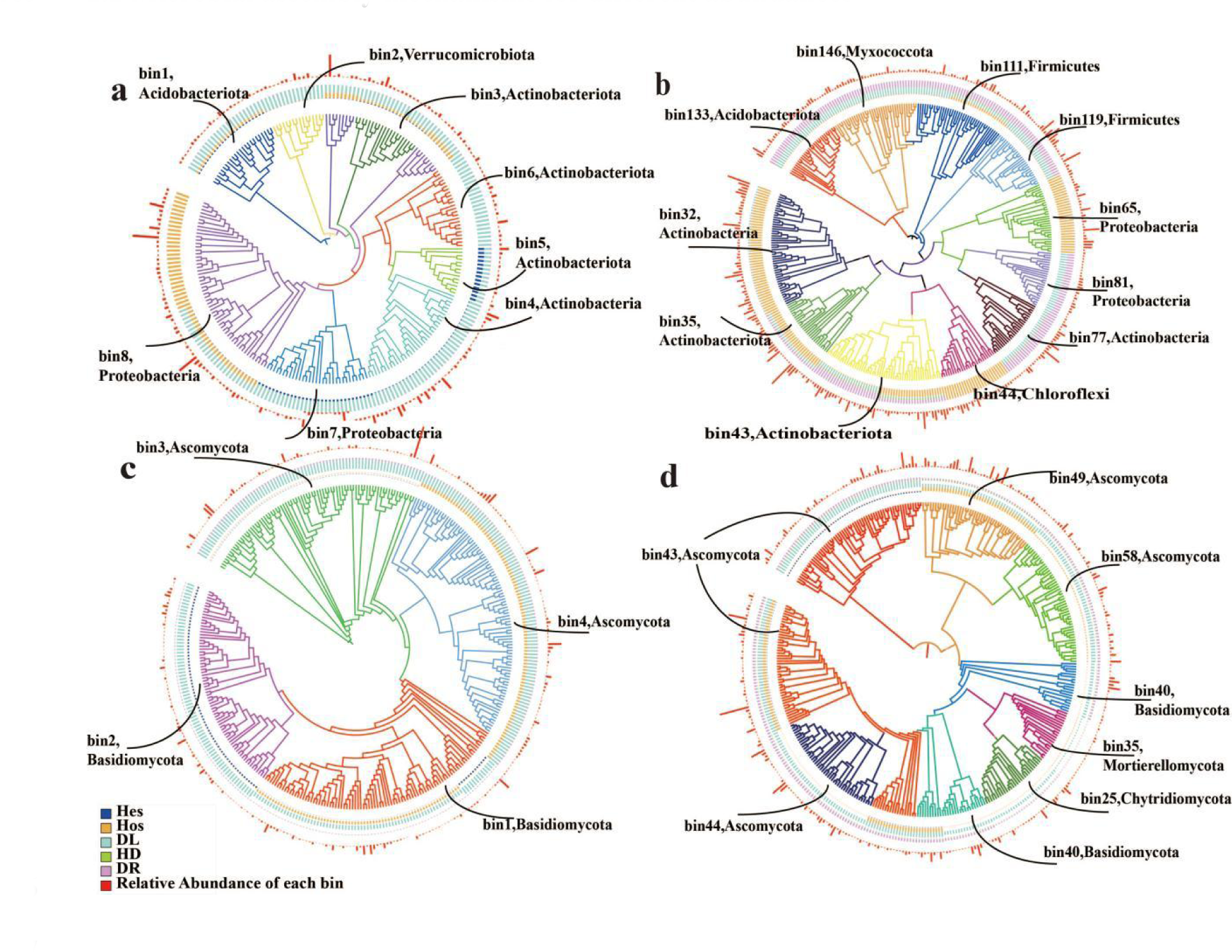
Mechanisms of assembly of microorganisms of different abundance (inner layer is the proportion of ecological processes, outer layer is the relative abundance of each bin; a: abundant bacteria, b: rare bacteria, c: abundant fungi, d: rare fungi)

Cluster tree results showed that among the rich microorganisms, soil samples from broadleaf forests were the most distant, followed by samples from mixed forests, with conifers, shrubs and herbs closer together. Rare microorganisms were more distant from each other than abundant microorganisms, showing closer proximity across samples in trees, followed by shrubs, with the most distant microorganisms in grassland samples. Fungal β-diversity of different abundances showed a consistent trend, all showing the greatest distance in mixed samples, followed by herbaceous, followed by pure forest.

### 2.2 Microbial assembly mechanisms for different abundances

In this study, the process of community assembly is divided into five components; HeS refers to the situation where a highly spatially heterogeneous environment reduces phylogenetic differences between communities, leading to a different community structure; HoS, on the contrary, refers to the situation where a spatially homogeneous environment gives rise to community homogeneity; while HD describes the diffusion between communities, which may occur in the soil through processes such as physical perturbation, water infiltration, or active diffusion in the aqueous film or in saturated pore space occurring in the soil; DL refers to spatial segregation; and DR describes ecological drift. Based on the iCAMP model, abundant and rare bacteria were classified into 41 and 160 phylogenetic bins, respectively, a number that has counterparts of 28 and 60 in fungi. Abundant subcommunities, both fungi and bacteria, were dominated by stochastic processes, and the order in different vegetation types was specifically presented as K > H > Z > C > G. Diffusion limitation (DL) was absolutely dominant in bacteria, and DL was dominant in fungi, followed by DR. Rare taxa had a higher proportion of stochastic processes, which were presented in specific types as K > C > G > H > Z, with 70% of the boxes being DL processes and 23% were DR processes.

For specific boxes, in the rich taxa, bin7 in bacteria is mainly controlled by HoS, bin5 is dominated by HeS in the forest samples, and DL is dominant in the other boxes, while in fungi DL is dominant except for bin4, which is HoS in the shrub samples. The situation is more complex in the rare sub-communities, where the dominant process in bacteria is deterministic, with the proportion of H > G > K > Z > C in all samples. Among all bins, the proportion of HoS (44%) is the highest, with DR (42.67%) being slightly lower, followed by DL (14%). Further comparing the major microbial taxa contained in the selected bins, the major bacterial taxa controlled by HoS and HD included *Aspergillus, Acidobacteriota, Actinobacteria, Verrucomicrobia* and Thick-walled Bacteria; and the major fungal taxa included *Aspergillus, Ascomycetes, and Spliceomycetes*. It is worth mentioning that the dominant microbial taxa controlled by HoS and HD belong mainly to rare taxa.

### 2.3 Soil factors drive microbial assembly processes

Environmental thresholds and phylogenetic signals of ecological preferences of abundant and rare taxa were tested under complex environmental gradients to reflect their environmental adaptations, and Figure 4 shows that the ecological niches of abundant taxa are wider than those of rare taxa, and that the ecological niche widths of bacterial taxa are wider than those of fungal taxa; abundant taxa exhibit stronger phylogenetic signals compared to rare taxa.

**Fig. 4.**
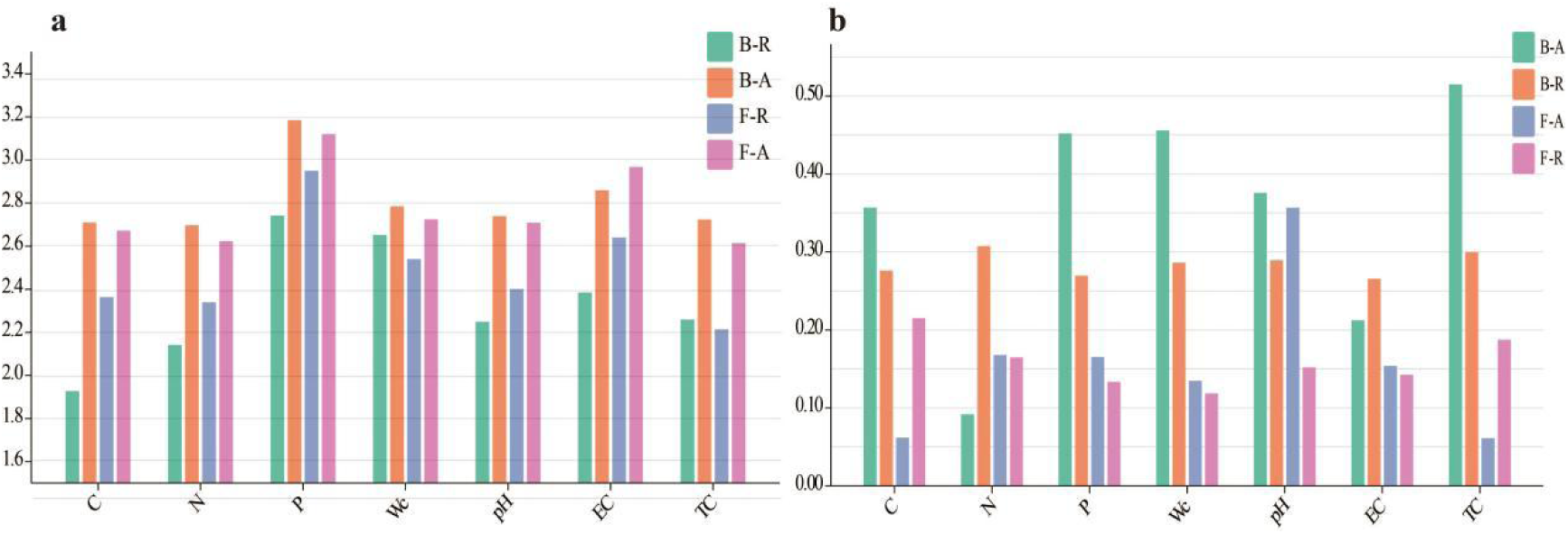
Ecological niche of soil environmental factors (a) and phylogenetic signals (b) for different microorganisms(B-R:abundant bacteria;B-A:rare bacteria;F-A:abundant fungi;F-R:rare fungi)

The heat map(Figure 5) further reveals the distribution patterns and assembly mechanisms of soil microorganisms of different abundances as influenced by soil factors. Rare taxa, both fungi and bacteria, were affected by more environmental factors. The NTI and Shannon index of abundant microorganisms were mainly affected by soil nutrient content while soil temperature had a significant effect on rare taxa; among the processes related to community building, HeS and HoS were mainly negatively correlated with soil factors while DL and HD showed mainly positive correlations, and regarding the DR process, except for rare fungi, which were negatively correlated with soil factors, it was positively correlated with soil factors for all other taxa. Specifically, the HeS process in the enriched taxa was mainly negatively correlated with nutrients and water content in the soil, with fungi (cor was −0.61) showing a more pronounced correlation than bacteria (cor was −0.37). The HOS process was controlled by soil nutrients. The DL process was similar to HD, and was mainly affected by CNP in bacteria whereas soil physical properties were more affected in fungi, e.g., by PH, water content, and soil temperature. Soil factors mainly influenced the DR process in abundant taxa and to a lesser extent in rare taxa.

**Figure 5.**
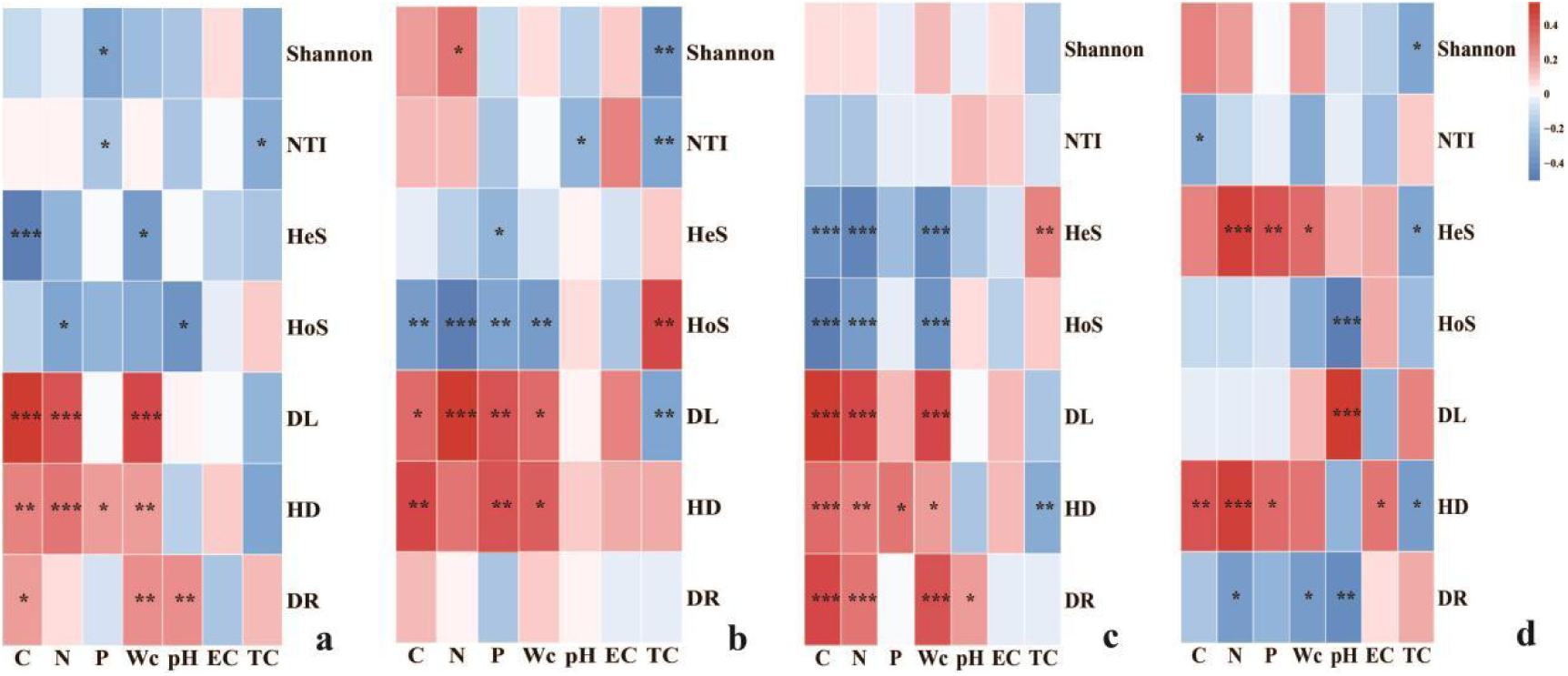
Correlation between the microbial assembly process and the environmental factors(a: abundant bacteria, b: rare bacteria, c: abundant fungi, d: rare fungi)

## 3. Discuss

### 3.1 Composition of microorganisms of different abundance

Rare and abundant microbial subcommunities have different compositions in different ecosystems. In this study, rare taxa dominated the majority of OTUs (98.14 % for bacteria and 95.12 % for fungi), but their sequences accounted for only 21.73 % (bacteria) and 33.78 % (fungi) of the entire community. In all the sample sites, both bacteria and fungi, the abundance of abundant taxa was higher than rare microorganisms but with fewer species, this is due to the fact that abundant taxa have easier access to nutrients than rare taxa, which leads to their greater abundance in the same habitat [12], whereas rare bacteria have a greater abundance of species, allowing a large number of specialists to adapt to the environment and thus efficiently utilise the resources of the wider range of woodlands. Regarding the composition of species at the genus level, the composition of bacteria and fungi is different, with rare and abundant bacterial species in bacteria differing slightly in composition, whereas the composition of fungi is more consistent. Specifically, rare bacterial species are mainly composed of oligotrophic taxa, such as *actinomycetes*, because rare micro-organisms are forced to exploit new survival resources with oligotrophic taxa as abundant species have already occupied optimal ecological niches; whereas fungi are able to access resources due to the presence of mycelium and other reasons for easy migration, so there is little difference in the composition of rare and abundant taxa.

The results of diversity index showed that rare taxa had higher α and β diversity and richness than the abundant taxa, indicating that rare taxa are more diverse and can adapt to changes in the environment. For different vegetation types, Shannon’s index (5.575) and observed species (331) were the highest in mixed forests, indicating that the diversity and richness of bacterial communities in this type were higher than other samples. An increase in *acidobacteria* in coniferous forests caused a decrease in the relative abundance of *actinobacteria*, as these taxa share similar ecological niches. Habitat heterogeneity was typically much higher in broadleaf and mixed forests compared to grassland samples due to the highly diverse environmental characteristics, resulting in significantly higher soil microbial richness. Herbs and shrubs were not clustered with other samples in the clustered tree-based beta diversity results, suggesting that the bacterial community in the scrub differed from that in the arboreal samples. Differences in abundance between sites can often be explained by the interaction of environmental selection and dispersal limitation on soil microbial abundance, and the higher beta diversity of rare taxa may indicate external microbial inputs by dispersal. Rare bacterial taxa showed greater variation in community similarity between types than abundant taxa, suggesting that rare bacterial communities are more sensitive to biotic factors. The greater the spatial variability within a microbial community, the more susceptible it is to environmental change, and the species composition of rare bacteria is more susceptible to geographic and environmental filters, resulting in much lower community similarity for rare bacteria than for abundant bacteria. Notably, the proportion of unclassified sequences was higher in rare taxa (10.2%) than in abundant taxa (2.6%), and rare taxa were more likely to become extinct or to fall below detection levels, all of which suggests that uncharacterised diversity is much greater in the rare biosphere.

### 3.2 Assembly processes in microorganisms of different abundance

Most of the previous studies were limited to community aggregation processes at the sub-community level, this study conducted in-depth studies of community building mechanisms at both the sub-community and phylogenetic bin levels, and the results showed that stochastic processes (e.g., diffusion limitation) contributed more to the composition of abundant sub-communities in the forest soils, whereas among rare microorganisms, the main controlling processes were variable selection for bacteria and diffusion limitation for fungi, which was in line with the previous finding that abundant taxa are more subject to environmental constraints [13]. The Icamp model results showed that diffusion limitation dominated the rich community assembly by 44-60% of the ecological processes and had the same results in all types. Rare taxa may be at a disadvantage in competition for limited resources, leading to the dominance of deterministic processes in rare taxa [14]. Abundant taxa, due to their higher abundance, are more susceptible to dispersal constraints, as opposed to rare taxa, which are local ‘experts’ with low abundance, limited dispersal capacity, and heterogeneous distribution across sites, thus are more susceptible to environmental filtering [15], and are less resistant to environmental change than abundant taxa occupying a wide range of ecological niches. Resistance to environmental change is not as strong as that of rich taxa occupying broad ecological niches. The contribution of drift and homogeneous diffusion to microbial subcommunity aggregation increased significantly with increasing rarity, while the contribution of homogeneous selection followed the opposite trend. According to probabilistic theory, subcommunities with higher rarity are themselves more susceptible to stochastic processes such as drift and diffusion than abundant subcommunities. Second, a large number of rare taxa are dormant in their environment; they are usually less active but more resistant to environmental changes [16]. Thus, rare subcommunities are less susceptible to deterministic processes such as selection.

While stochastic processes are important for bacteria, especially *Acidobacter* and *Actinomycetes*, soil fungi are usually distributed around roots, and thus mycelium formation leads to a more stable soil fungal community that is less likely to be affected by drift. The reason that fungi tend to show more diffusion limitation than bacteria in various ecosystems is that fungi are more likely to be limited in long-distance dispersal than smaller-sized bacteria, as the size of the organisms affects their dispersal capacity and spatial aggregation. The rate of spreading is highly dependent on species traits (flagella, cell size, resistance capacity, etc.) and the active state of the bacterial taxa [17]. The main propagules of fungi are spores or ascospores, which can be dispersed over long distances in the soil. Fungal dispersal is usually a passive process mediated by wind, water and animals [18], and is therefore susceptible to dispersal constraints, whereas most bacteria have their propagules as themselves, and are more susceptible to harsh abiotic conditions when dispersing over long distances than fungi [19]. However, at the same time, bacteria can also spread through fungal hyphae due to their small size, high abundance and rapid reproduction. As the role of dispersal limitation fades, the role of variable selection becomes apparent, and mixed forests increase the unpredictability of microbial composition and the stochastic nature of microbial assemblages through reduced resource competition, ecological niche selection, and enhanced preferential effects.

Comparing the different vegetation types, deterministic processes (heterogeneous selection) dominated the bacterial and fungal communities in coniferous forests (69.4% for bacteria; 88.9% for fungi), stochastic processes dominated the bacterial community in broadleaf forests (77.8%), and deterministic processes dominated the fungal community (52.8%), with mixed forests and shrub forests falling in between. This is due to the fact that stochastic processes dominate in high diversity communities, while deterministic processes dominate in low diversity communities [20]. It has been observed that the establishment of fungal communities in both coniferous and broadleaf forests in northern China is dominated by deterministic processes, and the formation of bacterial communities is dominated by deterministic and stochastic processes, respectively [21]. In this study, microbial assemblages in mixed forests with high diversity increased stochasticity through reduced resource competition, ecological niche selection, and enhanced preference effects, in contrast to fungal community aggregation under monocultures, which was dominated by stochastic processes, possibly due to the strong regional character of fungi and their limited dispersal [22]. This can be interpreted as a legacy effect of plant species, where the influence of tree species characteristics and long-term single-apomictic inputs leads to relatively specialised habits, which results in a greater disparity in environmental conditions between coniferous and broadleaved pure stands. Changes in heterogeneous environmental conditions increase turnover in community composition and, as a result, single stands increase heterogeneous selection and lead to high levels of variation in soil bacterial communities. There are three basic mechanisms that explain why stochastic processes are more influential in mixed stands. First, stochastic processes have a greater influence on community assembly when environmental factors do not exert an overwhelming selection pressure. Second, mixed stands have greater nutrient availability than pure stands, which may introduce more stochastic effects on microbial communities. Finally, because of the more homogeneous environment, diffusion processes may be higher in mixed forests than among monocultures, and may even outweigh the effects of deterministic processes at high diffusion rates.

### 3.3 Influence of soil environmental factors on microbial assembly processes at different abundances

Various environmental factors play a key role in driving the ecological assembly of microbial communities. Rare microorganisms show wider environmental breadth in all environmental parameters except soil temperature compared to enriched taxa [23], suggesting that rare species have narrower environmental thresholds, are less environmentally adapted, and are more susceptible to environmental selection, and Figure 4 illustrates that rare bacterial taxa comprise more specialists, whereas abundant taxa comprise more polymorphs. Abundant taxa are generally considered to be the most active classes in biogeochemical cycles, particularly carbohydrate metabolism, and occupy core ecological niches. The intrinsic genomic traits and wider ecological niches of enriched taxa allow them to be universally distributed with more individuals that are easily dispersed and can efficiently access a wider range of resources than rare taxa. The wide ecotope width and even distribution of the traits suggest that abundant taxa have greater metabolic versatility than rare taxa, largely due to their competitiveness in terms of resources and resistance to abiotic changes. The results of the heatmap(figure 5) further illustrate that the different assembly processes between rare and abundant bacterial communities may be due to differences in response to environmental disturbances and ecological niche breadth. Positive mean NTI for abundance taxa and both positive and negative results in rare communities suggest that the assembly of abundant taxa is dominated by phylogenetic clustering, with clustering and dispersal constraints controlling mainly the rare subcommunities. Higher NTI values can indicate not only stronger environmental filtering, but also greater genome diversification at the OTU level allowing for the coexistence of more closely related organisms due to higher mutation rates in populations of microbially evolved individuals under stressful conditions than in normal environments. According to Blomberg’s K-statistics, abundant microorganisms have stronger phylogenetic signals across all topographic and physicochemical parameters, and larger K values imply defined evolutionary processes, suggesting that abundant microorganisms may have had higher phylogenetic ecological niche conservatism, i.e., microbial adaptations to persistent environmental change, throughout their evolutionary history. The ecological niche results also support this conclusion: the larger the phylogenetic clustering of the community, the more likely it is that strong selection is occurring, because ecological traits that conserve ecological niche breadth are usually highly conserved. Without selection, communities may experience overdispersion, leading to more similar communities than would be expected by chance. These results suggest that rare and abundant microorganisms exhibit opposite environmental adaptations at the taxonomic and phylogenetic levels, while most soil factors do not directly affect soil bacterial similarity, but rather modify environmental selection pressures by increasing environmental similarity.

In this study, it was found that soil temperature, phosphorus content and vegetation type had highly significant effects on abundant microorganisms, while soil nitrogen content, soil moisture content, soil temperature and vegetation type had highly significant effects on rare microorganisms. Bacteria are more resilient than fungi because the filamentous nature of fungi makes them more sensitive to environmental changes at the micron scale. Previous studies have shown that soil pH [24] and moisture have been key factors influencing microbial community structure. pH is closely related to microbial community composition in woodland soils, altering microbial diversity through the effects of cell membrane structure and ion transporters. Tests of correlations with environmental factors (Fig. 5) also showed that significant positive correlations of NTI values of rare soil taxa with temperature and pH imply that the combination of these abiotic factors exerts greater pressure on the communities within them, and that selective pressures result in close taxonomic correlations (i.e. higher NTI), but that environmental heterogeneity and variations in environmental conditions increase taxonomic distinctiveness (i.e. higher diversity). The decline in the relative importance of dispersal constraints to sub-communities of rare fungi may have led to diminished differentiation. Bacteria require wider ecotope widths increasing the likelihood of successful colonisation when spreading to new locations. Dispersal limitation implies that low rates of dispersal are the main cause of different community structures, where drift (stochastic changes in species abundance) becomes more important for community composition due to low turnover and weak selection. Moist habitats promote microbial dispersal. The decisive role of soil electrical conductivity (EC) in influencing the assembly of rare and abundant bacterial taxa may be due to the toxicity of EC on the growth activity of microbial cells. Low EC favours microbial metabolic activity and growth, which can increase microbial dispersal capacity. High EC may lead to greater microbial-to-microbial rejection and reduced colonisation, leading to a greater impact of deterministic assembly on the microbial community [25]. Notably, a significant effect of soil temperature on microbial community assembly was observed in this study. This can be attributed to temperature affecting the metabolic rate of microorganisms and their competitive interactions [26] or enhancing the solubilisation of mineral elements and accelerating the rate of redox reactions affecting resource availability [27], leading to changes in microbial community assembly.

In addition, soil nutrient content is also an important factor, and the resource availability-stochasticity relationship proposed by Dini-Andreote, de Cássia Pereira e Silva [28] showed that high resource availability can increase stochasticity in the absence of strongly selective physico-chemical conditions, with deterministic processes tending to occur under low-nutrient and harsh conditions, and stochastic processes occurring under nutrient-rich and mild conditions. This supports the result that the greater proportion of stochastic processes in mixed forests (Figure 3) is due to the fact that nutrient abundance reduces competition between microbial communities, which explains the greater proportion of stochastic processes in forests than it does in grasslands. Increasing soil resource availability reduces deterministic processes, and plants in the community act as the main biofilter for soil microbes and can provide many specialised ecological niches for rare taxa by diversifying the substrate of dying leaves and root exudates and by forming symbioses, which alter microbial assembly processes. Rare bacterial taxa are more susceptible to soil nutrient variability than abundant bacterial taxa, possibly because changes in soil nutrients may drive bacterial communities towards phylogenetic dispersal, causing fluctuations in rare taxa. Deterministic selection and dispersal limitation were mainly mediated by soil effective phosphorus, and this influential process was stronger in fungal than in bacterial communities. Soil phosphorus increases the total soil carbon content by promoting plant growth, thus altering the community structure. On the other hand, phosphorus can also influence microbial growth by altering soil pH and osmotic pressure [29], while the availability of nitrogen and phosphorus shapes fungal communities by affecting fungal enzyme activities and functional gene abundance. In phosphorus limited soils, legume roots mainly secrete citric, fumaric, malonic, succinic and malic acids, which favours the growth of Aspergillus, whereas in phosphorus rich soils stimulates the growth of Acidobacteria [30], which explains the presence of large amounts of Acidobacteria syncytialis Aspergillus under the herbaceous sample plots, which contained a wide range of leguminous species (Figure 2).

## 4. Materials and Methods

### 4.1 Study area

The Tian Shan mountain range is located in Central Asia, and most of it lies within the Xinjiang Uygur Autonomous Region of China, which has a temperate continental climate, with an average annual precipitation of 1200 mm, an average annual temperature of 2-3°C, and a rich variety of vegetation species with a high cover, presenting a non-zonal character. In mid-July 2022, soil/leaf samples were collected from five forests and grasslands (Coniferous forests: Z, broadleaved forests: K; mixed forests: H; shrub forests: G; herbs as control: C) in three typical montane forested areas selected on the northern slopes of the Tianshan Mountains with longitude variation. The specific study areas were located in Barkun Forest (DTS), Urumqi Forest Parkt (ZTS) and Wusu Forestt (XTS), Xinjiang, China, respectively, with grey-brown soil type in the understory, herbaceous plant communities dominated by *Taraxacum mongolicum*, *Geranium wilfordii*, and *Potentilla chinensis*, and shrubby plant communities dominated by *Rosa platyacantha*, *Caragana sinica*, and *Juniperus pseudosabina*.

### 4.2 Sample collection

At each of the three sites, three sampling replicates of each type were selected to exclude anthropogenic interference, and the distance between adjacent plots was greater than 100 m. Soil was sampled at five points to remove visible litter, stones, etc., and a total of 45 soil samples were obtained, which were later divided into three sub-samples that were preserved at −80°C, 4°C, and room temperature (air-dried), respectively. In addition, we recorded the longitude and latitude of the sampling sites using GPS. The plant cover and number of species in each sample was also recorded.

### 4.3 Determination of physical and chemical properties of soil

The soil was dried at 105°C for 48 h to determine the moisture content, the pH of the soil was measured by a pH meter, the EC of the soil was determined by a conductivity meter, the carbon content of the soil was determined by the oxidation method of potassium dichromate, and the content of total phosphorus and nitrogen of the soil was determined by the molybdenum-antimony spectrophotometric method (HJ 632-2011) and the Kjeldahl nitrogen determination method (HJ 717-2014), respectively.

### 4.4 DNA extraction and NovaSeq sequencing

Soil DNA was extracted using the DNeasy PowerSoil ® kit (Qiagen, Hilden, Germany) according to the manufacturer’s instructions. The PCR run started with an initial denaturation and enzyme activation step at 95°C for 10 min, followed by 40 cycles of 30 s at 95°C, 30 s at 56°C and 45 s at 72°C, with an extension of 72°C for 10 min. PCR products were purified using the GeneJET Gel Extraction Kit (Thermo Scientific, Lithuania) and subjected to Illumina NovaSeq High-Throughput Sequencing using a double-ended strategy with a sequencing depth of over 30,000 sequences per sample.

### 4.5 Construction of the phylogenetic tree

All Effective Tags of all samples were clustered using the Uparse algorithm (Uparse v7.0.1001, http://www.drive5.com/uparse/), and sequences were clustered into OTUs (Operational Taxonomic Units) with 97% identify. Species annotation was performed on the OTUs sequences, and the phylogenetic relationships of all OTUs representative sequences were obtained by rapid multiple sequence comparison using MUSCLE (http://www.drive5.com/muscle/) software. For bacterial communities, phylogenetic trees based on 16S rRNA gene sequences were constructed in FastTree software using the “maximum likelihood” method and Figtree software. For fungal communities, phylogenetic trees based on ITS sequences were constructed in Ghost-tree software.

### 4.6 Data analysis

Differences in microbial abundance and diversity between vegetation types and study sites were analysed based on ANOVA and significance tests. Relationships between soil microbial diversity and phylogenetic indicators and soil properties were determined by Pearson correlation analysis. Habitat ecotope widths of microbial taxa are key factors to consider when studying deterministic and stochastic processes, the TITAN2 package was used to calculate ecotope widths (Levins’ ecotope widths) for different microbial taxa. Analysed the relative impacts of different ecological processes using the iCAMP package in R4.3.1.

### Data availability

The data used in this paper are high-throughput sequencing data, stored in National Center for Biotechnology Information, PRJNA989514.

## 5. Conclusions

Using phylogenetic bin-based null model analysis (iCAMP), this study explored the processes and their main drivers affecting the assembly of soil bacterial and fungal communities under different vegetation types. The results showed that (1) rare taxa of microorganisms exhibited higher diversity and stronger phylogenetic clustering, and enriched microorganisms had a wider width of environmental ecological niches; (2) variable selection and diffusion limitation proved to be the most important assembly processes for bacterial and fungal communities, respectively; (3) soil temperature and nutrient effectiveness mediated the relative importance of deterministic and stochastic processes in soil microbial communities. This finding provides valuable insights into the microbial diversity and assembly processes in forest soils in the arid zone of northwestern China and reveals the extent to which community assembly processes are uniquely influenced by the soil environment.

## Acknowledgments

This work was supported by The third comprehensive scientific investigation project in Xinjiang(2021XJKK0900)and Key Laboratory of Oasis Ecology.

